# Connecting Syncmers to FracMinHash: similarities and advantages

**DOI:** 10.1101/2023.11.09.566463

**Authors:** Shaopeng Liu, David Koslicki

## Abstract

**Motivation:** Sketching methods provide scalable solutions for analyzing rapidly growing genomic data. A recent innovation in sketching methods, syncmers, has proven effective and has been employed for read alignment. Syncmers share fundamental features with the FracMinHash technique, a recent modification of the popular MinHash algorithm for set similarity estimation between sets of different sizes. Although previous researchers have demonstrated the effectiveness of syncmers in read alignment, their potential for broader usages in metagenomic analysis (the primary purpose for which FracMinHash was designed) and sequence comparisons remains underexplored.

**Results:** We demonstrated that a open syncmer sketch is equivalent to a FracMinHash sketch when appled to *k*-mer-based similarities, yet it exhibits superior distance distribution and genomic conservation. Moreover, we expanded the concept of *k*-mer truncation to open syncmers, creating multi-resolution open syncmers for metagenomic applications as well as flexible-sized seeding for sequence comparisons.

**Reproducibility:** All analysis scripts can be found on [GitHub](https://github.com/KoslickiLab/FMH_vs_syncmer_reproducible).

## Introduction

*K*-mer-based methods are widely used in bioinformatics analyses, particularly in metagenomics. Sketching methods, which involve selecting a subset of distinct *k*-mers (referred to as a “sketch”) generated from genomic data, have become indispensable, enabling efficient computational comparisons for large-scale datasets [15, 17, 11]. Two notable methods in this category, FracMinHash [10] and Syncmers [5], are both designed to generate random samples (sketches) of all *k*-kmers from arbitrary genomic sequences.

FracMinHash, an extension of the popular MinHash algorithm tailored to estimate set similarity, particularly when set sizes vary significantly, selects a fixed portion of *k*-mers by applying a proportional cutoff to the hash space to which *k*-mers are randomly projected [10]. FracMinHash has gained widespread adoption in metagenomics for conducting extensive genomic comparisons such as taxonomic profiling [17] and ANI estimation [21, 12]. On the other hand, Syncmers select *k*-mers characterized by the presence of their smallest substring at specific location(s) [5]. For example, open syncmers refer to *k*-mers whose smallest substring of length *s* shows up at their *t*-th location. Notably, Syncmers have found successful applications in read aligners, surpassing the widely-used Minimizer method in performance [20, 4, 18, 19].

It’s important to note that these two methods operate in different contexts. In metagenomics, alignment-free genomic comparisons are usually performed using distinct *k*-mer sets, though there are algorithms based on weighted *k*-mers, generated from genomic sequences. On the other hand, in read alignment tools with the seed-and-extension algorithm, the duplication of *k*-mers is a consideration for seed matching status and an important consideration for optimizing speed and time [20]. Unless explicitly stated, we will primarily focus on the former scenario, as the latter has already been extensively discussed.

These two methods have three core attributes in common: 1) Context-free (1-Locality); the selection of a *k*-mer is solely determined by its own content and is not influenced by adjacent bases. Therefore, if a particular *k*-mer is chosen from one sequence, it will also be selected from other sequences. 2) random and fractional sample (sketch); both methods randomly select *k*-mers with sample sizes proportional to the original *k*-mer set size. 3) *k*-mers in each sketch are min-wise independent (discussed in section 2.2.3), allowing them to estimate the Jaccard and containment indices.

Conversely, the primary distinction between them lies in the situation of overlapping *k*-mers. In FracMinHash, hash values of every *k*-mer are independent and identically distributed. Therefore, the events of being selected into a FracMinHash sketch are independent of overlapping (i.e., *k*-mers sharing substrings are independent of being selected into a FracMinHash sketch). In Syncmers, however, overlapping *k*-mers can influence one another in terms of their likelihood of being classified as Syncmers (Fig 1). While syncmers have demonstrated their effectiveness in read alignment, their potential for use in genomic analysis has not been fully realized.

**Fig. 1.**
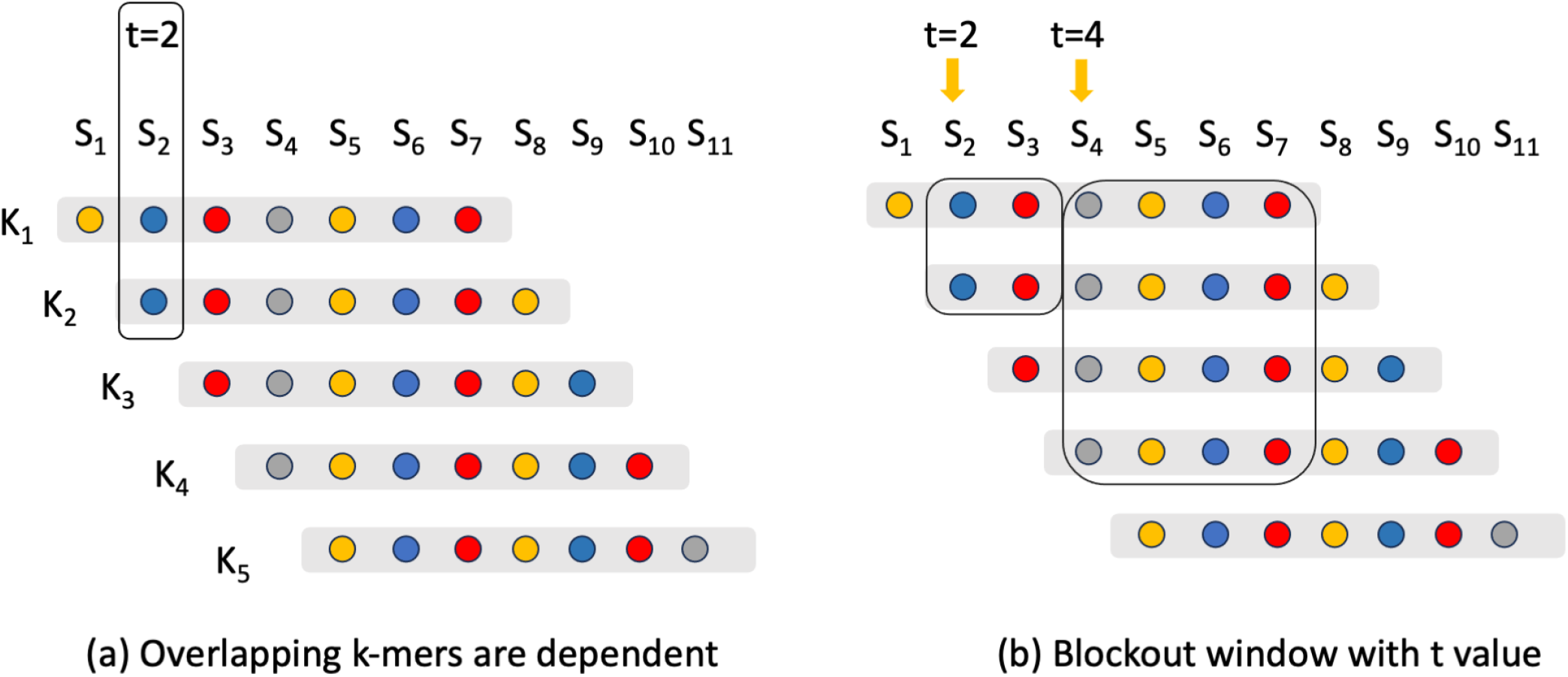
Overlapping *k*-mers are dependent on the status of being an open syncmer. There are 5 consecutive *k*-mers and each *k*-mer has 7 substrings (every dot represents a substring of length *s*). (a) An open syncmer may affect its overlapping *k*-mers regarding the likelihood of being an open syncmer. When *t* = 2, if *K*_1_ is an open syncmer (i.e., *S*_2_ *< S*_3_), then *K*_2_ must not be an open syncmer. (b) When *t >* 1, every open syncmer will have a blockout window where some downstream *k*-mers must not be an open syncmer. If *K*_1_ is an open syncmer: when *t* = 2, we have *S*_2_ *< S*_3_ so *K*_2_ must not be an open syncmer; when *t* = 4, we have *S*_4_ *< S*_5_, *S*_6_, *S*_7_ so *K*_2_, *K*_3_, *K*_4_ must not be an open syncmer.

**Fig. 2.**
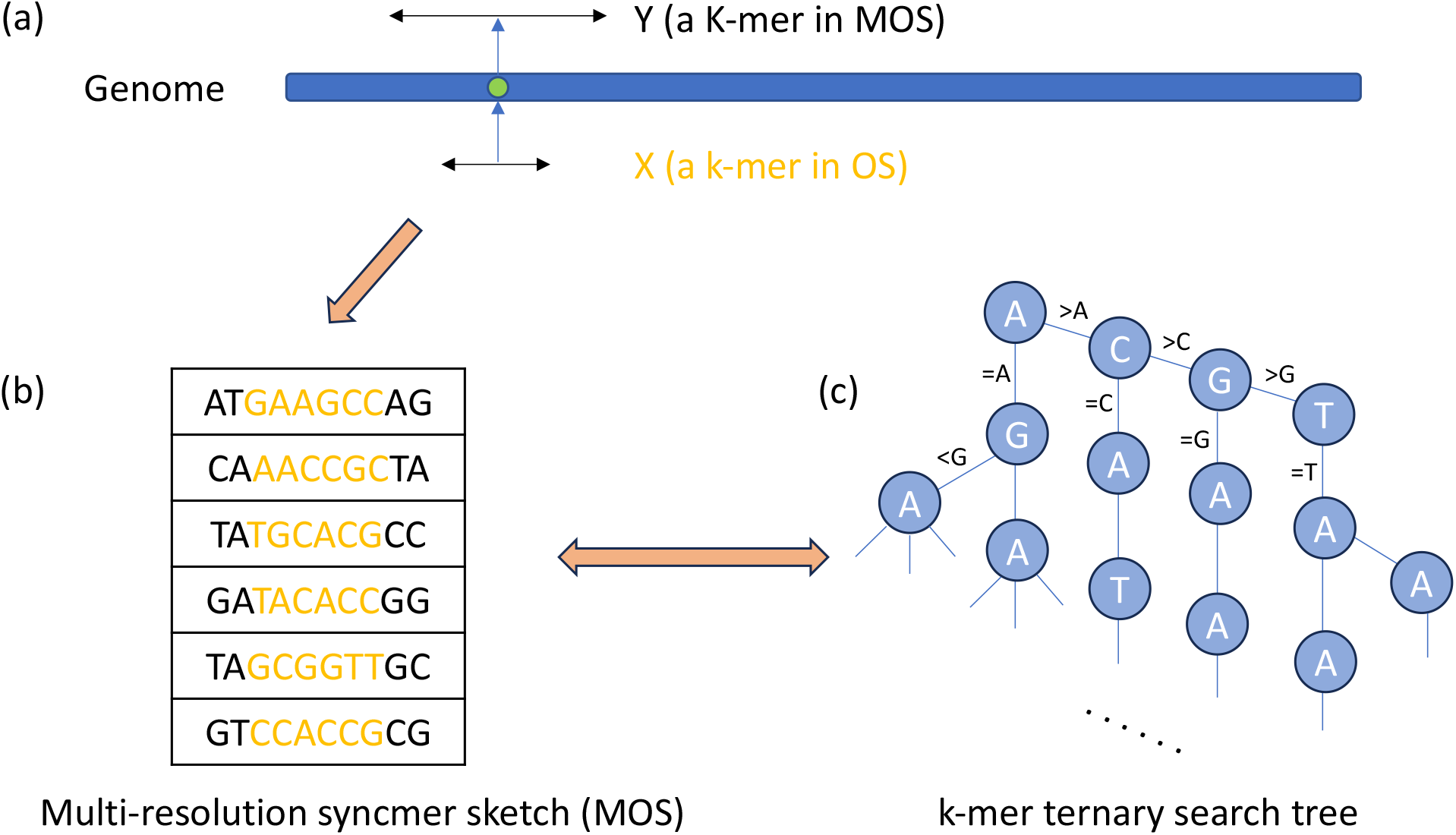
Multi-resolution open syncmer sketch. (a) Every *k*-mer from a syncmer sketch can be extended to length *K* to form a multi-resolution sketch (MOS). (b) A MOS by design contains the original OS sketch. (c) And it can be stored in a *k*-mer ternary search tree (or similar prefix trie) to enable prefix lookups to efficiently retrieve sketches for arbitrary lengths between *k* and *K*.

In this study, we conduct a theoretical analysis between open syncmers and FracMinHash sketches, demonstrating that open syncmers possess a similar capability to FracMinHash in performing MinHash-based similarity estimations. We validated our theoretical findings through empirical tests on microbial genomes. Meanwhile, we provided a theoretical foundation for open syncerms superior performance in genome conservation. Consequently, we argue that open syncmers are capable of conducting metagenomic comparisons on par with FracMinHash. Last, as illustrated in our previous research on multi-resolution *k*-mer-based estimations [14], the idea of *k*-mer truncation can be applied to open syncmers as well to fit various sizes to accommodate both sequence comparison and seed matching, indicating its potential for both read alignment and metagenomics analysis.

## Methods

### Preliminaries

#### Jaccard and containment index

In the metagenomics context, genomic comparisons are usually carried out using distinct *k*-mer (all substrings of length *k* extracted from genomic sequences) sets derived from the samples, with consideration given to the weight of *k*-mers (e.g. for quantification). Similarity between pairs of genomic data can be measured by the similarity of their respective *k*-mer sets: the collection of all distinct *k*-mers appearing as contiguous substrings in the data.

The Jaccard index (JI) measures the similarity of two non-empty sets by comparing the relative size of the intersection over the union and the containment index (CI) measures the relative size of the intersection over the size of one set [3]. We use *A*^*k*^ to denote the set of all distinct *k*-mers derived from a set of sequences/strings *A*. When applied to sets of *k*-mers, JI and CI are defined as follows:

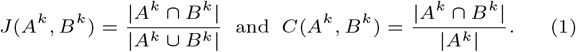

Typically, the value of *k* used is not small (e.g., *k*=21 for metagenomic profiling [17, 23]) such that it’s unlikely to have duplicate *k*-mers purely by chance. Nonetheless, real genomes are not random and contain repetitive regions. While duplicate *k*-mers are frequently encountered in practice, it is generally assumed that their occurrences are limited in number.

#### FracMinHash

The FracMinHash (FMH) sketching technique (a.k.a. mincode submer by Edgar [5], scaled MinHash by Irber et al. [10], and universe minimizer by Ekim et al. [6]) is a generalization of modulo hash that supports Jaccard and containment estimation between sets of different sizes.

In the analysis of real data, the MinHash algorithm employs a bottom sketch strategy [3]: given a sketch size *m*, a single hash function is used, instead of a family of *m* min-wise independent hash functions, to select elements associated with the smallest *m* hash values. FMH follows the same idea but samples a fixed proportion of *k*-mers [10].

Let [0, *H*] be the hash space of some perfect hash function *h* that can randomly project *k*-mers into the hash space, for a pre-selected scaling factor *c* (i.e., reciprocal of the proportion), a FMH sketch for sequences *A* is:

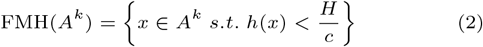

Therefore, it’s expected that 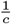 of all distinct *k*-mers will be selected at random, and the scaling factor *c* can be tuned to modify the sketch size. Besides, sampled *k*-mers are uniformly distributed along the genome (assuming no repeat regions) as every *k*-mer has an equal probability of being selected along the genome.

#### Open syncmers

Open syncmers (OS) represent a *k*-mer sampling approach involving the random selection of *k*-mers based on the relative order of their substrings. Open syncmers are defined by a *triple* (*k, s, t*): a *k*-mer becomes an open syncmer if its smallest (relative to some hash function) substring of length *s* (*s < k*) shows up at a specific location *t*. Using (*k* = 5, *s* = 3, *t* = 1) as an example, a 5-mer *AAGCT* is an open syncmer because it has 3 substrings *h*(*AAG*) *< h*(*AGC*) *< h*(*GCT*) where the 1st substring is the smallest if the *h* simply reflects the lexicographical order. An open syncmer sketch for sequences *A* is:

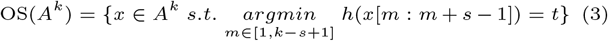

Intuitively, a *k*-mer will have (*k* − *s* + 1) substrings of random order. It will be selected as an open syncmer with a probability of 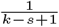. This setting is similar to the FracMinHash technique except for the dependence on overlapping *k*-mers: they are not independent regarding their classification as open syncmers because they share substrings (Fig 1).

Various types of syncmers exist [5]. We opted for open syncmers because they already offer an excellent genome compression ratio and genome conservation (ratio of bases covered by matching *k*-mers between a genome and its mutated sequences); and they have demonstrated successful applications in minimap2 [13] and Strobealign [18]. A generalized approach, parameterized syncmer scheme [4], has been discussed to include an arbitrary number of locations for the smallest substring, with open syncmers representing the case of only one substring location.

Syncmers can be further downsampled to accommodate a larger compression factor by employing the FracMinHash technique (referred to as “Mincode submer” in the original manuscript) [5]. Using a downsampling ratio *d* (*d* ≥ 1), a random *k*-mer will be selected as an open syncmer with a probability of 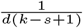. While keeping in mind that open syncmers can adapt to arbitrary scaling factors with followup downsampling, we assume *d* = 1 (no downsampling) for the following discussion without loss of generality.

Next, we will demonstrate, both theoretically and through practical examples, how an open syncmer sketch outperforms a FracMinHash sketch for genomic comparisons.

### Compare the open syncmer sketch to the FracMinHash sketch

In this section, we will compare the open syncmer sketch to the FracMinHash sketch from four perspectives: compression factor, *k*-mer distances, compatibility with the MinHash algorithm, and multi-resolution estimation.

#### Compression factor can be identical for both sketches

The compression factor (a.k.a. scaling factor) is calculated as the ratio of all distinct *k*-mers to the number of selected *k*-mers in a sketch of a long random string. Following the same argument as Edgar [5], it’s easy to deduce that both FracMinHash and open syncmers share the same expected compression factors by linearity of expectation no matter what the shift value *t* is. And we demonstrated this result in real data in section 3.1.

#### The open syncmer sketch provides better k-mer distances than the FracMinHash sketch

A subsampling method should avoid selecting *k*-mers with extensive overlaps or clustering (on genomes), as such *k*-mers provide less information compared to more widely distributed *k*-mers [8].

We can use a discrete random variable, denoted as *D*, to measure the distances between the starting index of two consecutive *k*-mers along the genome within a given *k*-mer sketch.

**Corollary 2.1**. *Both sketches have the same expected k-mer distance when c* = *k* − *s* + 1.

*Proof* When *c* = *k* − *s* + 1, the expected sketch sizes for both methods are the same. Considering a long random string *A* with *n* distinct *k*-mers in total and assuming a low repeat level,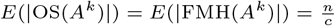.

The total distances between all adjacent *k*-mers in the sequence sum up approximately to the entire genome length, except a few preceding and trailing bases. Consequently, the average distances between two adjacent k-mers for both sketches are equivalent:

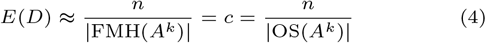

In practice, however, the average distances for both sketches are usually smaller than the expected value due to repeated regions in the genome. In the FracMinHash algorithm, the event that a *k*-mer is selected into the sketch can be treated as a Bernoulli trial with a probability of 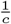 and these events are independent to each other. Therefore, we can derive that *D*_*F MH*_ follows a geometric distribution with success probability 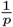 because the distance between two *k*-mers is the number of failures until the one event. Conversely, the intricate interdependence between overlapping k-mers for open syncmers (Fig 1) complicates the derivation of *k*-mer distances within an open syncmer sketch.

Interestingly, this interrelatedness plays a favorable role in open syncmers, distinguishing it from FMH. It effectively ensures that *k*-mers maintain a certain distance from each other, preventing overcrowding and thereby contributing to the desired properties of open syncmers.

### Open syncmers have lower bound for distances between adjacent *k*-mers

In the context of open syncmers with parameters (*k, s, t >* 1), every open syncmer creates a blocking window such that the next several consecutive *k*-mers must not be open syncmers (Fig 1b). This blocking event occurs when the *t*-th substrings of both the open syncmer (referred to as *K*_1_) and the other *k*-mer (referred to as *K*_*x*_) are covered in their overlaps. Using *K*[*i*] to denote the i-th substring of length *s* in a *k*-mer, the fact that *K*_1_ is an open syncmer implies that *K*_1_[*t*] *< K*_*x*_[*t*], making it impossible for *K*_*x*_ to be an open syncmer. The size of this blocking window is bounded by its distance from the prefix and suffix of a *k*-mer, as exceeding this distance would leave one of the *t*-th substrings uncovered by the overlapping segment. Consequently, the blocking window is symmetrical and achieves its maximum length when the *t* value is positioned at the midpoint of the *k*-mer. Namely, 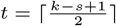 gives the optimal situation where every 2 consecutive open syncmers must be at least *t* bases away.

This observation aligns with the conclusion made by Shaw and Yu in [20, Theorem 8] that the optimal choice of *t* value is the middle of the *k*-mer and this gives the best genome conservation. Please note that selecting the optimal *t* value, while it improves base-level conservation with more scattered *k*-mers, does not impact sketch size or *k*-mer-level conservation (Figure 5b).

Syncmers with parameter *t* = 1 have the feature of window guarantee, a situation where a *k*-mer must be selected within a window of some length. But it has less practical relevance as the maximum possible distance can be large [5]. Therefore, we can derive the following based on the observation above:

#### Corollary 2.2.

*K-mers selected by open syncmer sketches in the genome are guaranteed to be t bases apart* 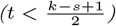 *assuming no repeats in the genome*.

Consequently, the distances between adjacent *k*-mers in syncmer sketches adheres to a lower bound (Corolloary 2.2) while maintaining the same expected value (Corolloary 2.1). This results in a more uniform distribution of *k*-mers along the genome and explains the better genome conservation in Fig 5.

#### The open syncmer sketch is suitable for the MinHash algorithm for full k-mer sets

As highlighted earlier, the MinHash and FracMinHash algorithms both utilize a bottom sketch strategy: they use a single hash function to uniformly and randomly project arbitrary *k*-mers into hash space to select elements by hash values instead of using a family of min-wise independent hash functions to select elements separately [3]. In metagenomic analysis, the single hash manner is mostly used for computational efficiency, and hash functions from murmurhash [1] family are popular choices.

Although not explicitly stated, it’s crucial to note that this randomness of hash projection is unrelated to the value of *k*. In other words, the hash values of different *k*-mers are independent, whether they share some substrings (overlapping) or one is a substring of the other. This property is known as the “avalanche property”, which characterizes a scenario where a minor alteration in the input causes a substantial alteration in the output, making it statistically indistinguishable from randomness (the widely used murmurhash family are such hash functions with excellent avalanche properties) [7].

In brief, under proper hash function *h* and a large hash space *H*, genomic sequences of arbitrary lengths are uniformly and randomly projected into the hash space. i.e.,

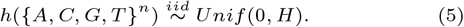

With this approximation, *k*-mers within the open syncmer sketch are uniformly and randomly distributed in the hash space. This is because each *k*-mer is selected based on the hash values of its substrings, which are unrelated to the hash value of the original *k*-mer. Restating these observations as a theorem, we have:

##### Theorem 1

*With a proper hash function h, k-mers in the open syncmer sketch are uniformly and randomly distributed in the hash space*.

Remember that both sketching methods choose *k*-mers based solely on their own *k*-mer sequences (1-locality), ensuring that if a specific *k*-mer is selected from one sequence, it will also be chosen from other sequences. Define *h*_*min*_(*S*) to be the minimal member of set S with respect to some hash function *h*, the probability that any *k*-mer in the open syncmer sketch possesses the minimum hash value is equal for all *k*-mers. That is to say, we can use the bottom sketch strategy to open syncmers to estimate OS-based Jaccard (*J*_*OS*_) and containment (*C*_*OS*_) indices.

**Corollary 2.3**.

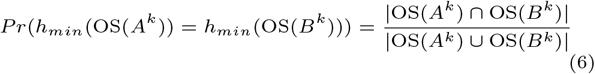

To be noted, minimizers can also be used for such computations but would give biased JI and CI estimation compared to the original k-mer sets [2]. Shibuya et al. demonstrated that closed syncmers (another format of syncmers) are comparable to random sampling and can provide reliable estimations for Jaccard Indices because closed syncmers are not context-dependent (1-locality) [22, Figure 1, Section 3.3]. Open syncmers follow the same logic and can be used for estimating regular JI and CI. To be specific, the theoretical work by Hera et al. [9, Theorem 1, Supplementary A.3] can be applied to open syncmers to show its efficacy in estimating *k*-mer set similarities, given that both open syncmers and FracMinHash sketches are fractional and random samples from the original *k*-mer sets.

### Estimate Jaccard and containment indices

Open syncmer sketches can be used to estimate Jaccard and containment index, just as MinHash sketches can. We then compare it to the FracMinHash sketch for estimating set similarity. Following the definition of FracMinHash given by Irber et al (2022) [10], we can get the estimate of Jaccard and containment indices via the FracMinHash sketch:

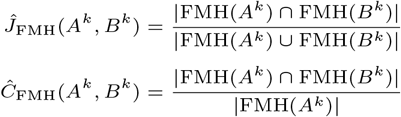

Similarly, the estimation by the open syncmer sketch:

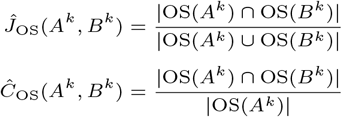

Please keep in mind that these estimations are not unbiased [9, Theorem 1, Supplementary A.3], but we may ignore the error term in practice as it is exponentially small with respect to the set sizes (which are appreciably large).

### Multi-resolution application with the open syncmer sketch

In our prior work CMash [14], we illustrated that the MinHash sketch is capable of offering multi-resolution estimations of Jaccard and containment indices by truncating a longer *k*-mer into shorter lengths. Now we broaden the *k*-mer truncation idea to include open syncmers for multi-resolution applications, encompassing both similarity estimation and seed matching. As previously discussed, we use 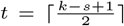 for open syncmers for the optimal genome conservation.

A multi-resolution open syncmer sketch (MOS) is defined by a regular open syncmer sketch (OS) with parameter 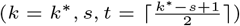 plus an additional parameter *K* (*K > k*^*^). A substring of length *K* is selected by MOS if it has an open syncmer of length *k*^*^ at its central position (any position works but the center is more convenient). Figure 2 gives an abstract idea of MOS: a MOS can be built upon a regular open syncmer sketch and can be further tuned to fit different substring lengths. This extension process preserves the advantages of syncmers, such as fractional sketching, high genome conservation, and inherent randomness, while also introducing versatility in adjusting the values of *k*. A formal definition would be:

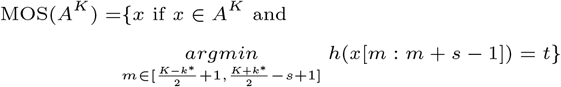

Then, MOS can facilitate *k*-mer-based applications with any *k* values between [*k*^*^, *K*] by simply truncating itself, eliminating the necessity to repeat the sketching/seeding step. When *K* = *k*^*^, equation **??** is exactly the definition of open syncmers and MOS is just the regular open syncmer sketch. In other cases, we can get an open syncmer sketch with new but smaller *k* sizes (preferably the difference is even) by simply truncating the MOS until length *k*.

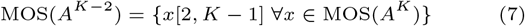

Coupled with a *k*-mer ternary search tree, we can store a single sketch for use across a range of *k* values, and conveniently modify the *k* size through prefix lookups. In addition to enabling multi-resolution estimations of *k*-mer-based similarities, the multi-resolution open syncmer sketch (MOS) offers a means to balance mapping quality and speed when utilizing a sketching based seeding approach in a seed-and-extend alignment algorithm. As demonstrated by Shaw and Yu, the use of open syncmers in minimap2 increases sensitivity but can lead to more repetitive seed matches [20]. Implementing multi-resolution seed matching has the potential to reduce repetition while preserving the sensitivity of smaller *k* values. In section 3.4, we will demonstrate the efficacy of applying MOS in metagenomic analysis.

## Results

In this section, we assess the performance of the open syncmer sketch in comparison to the FracMinHash sketch in a broad metagenomic setting. We obtained 20 random genomes in the Brucella genus (because these genomes exhibit a wide spectrum of Jaccard Indices) and 400 genomes from 10 randomly selected families from the GTDB database [16] to serve as reference sequences.

### Compression factor and *k*-mer conservation are consistent across open syncmers and FracMinHash sketches

The expected sketch size can be made equal if we choose parameters in such a way that *c* = *k* − *s* + 1. In our comparison, we set *k* = 20 and *s* = 11 for the open syncmer sketch and *c* = 10 for the FracMinHash sketch, resulting in both having an expected compression ratio of 10. We count via brute-force all distinct *k*-mers across all genomes and compared them to the sketches we generated. Similarly, *k*-mer level conservation (measured by the ratio of *k*-mers recovered from a mutated genome with 0.05 uniform random mutation rate) is consistent between FracMinHash sketches and open syncmers with different *t* values.

The results are presented in Figure 3, revealing that the compression factors align with our expectations and remain consistent irrespective of the value of *t*. Since open syncmers comprise a random selection from all distinct *k*-mers, we can anticipate that *k*-mer conservation will be similar to what we obtain from FracMinHash. As discussed previously, adjusting the value of *t* only affects genome conservation, not *k*-mer conservation. The *k*-mer conservation rate slightly exceeds the anticipated value, with the probability of a 20-mer remaining unmutated being 35.6%. This pattern persisted across various mutation rates, likely attributable to repetitive regions within the genome.

**Fig. 3.**
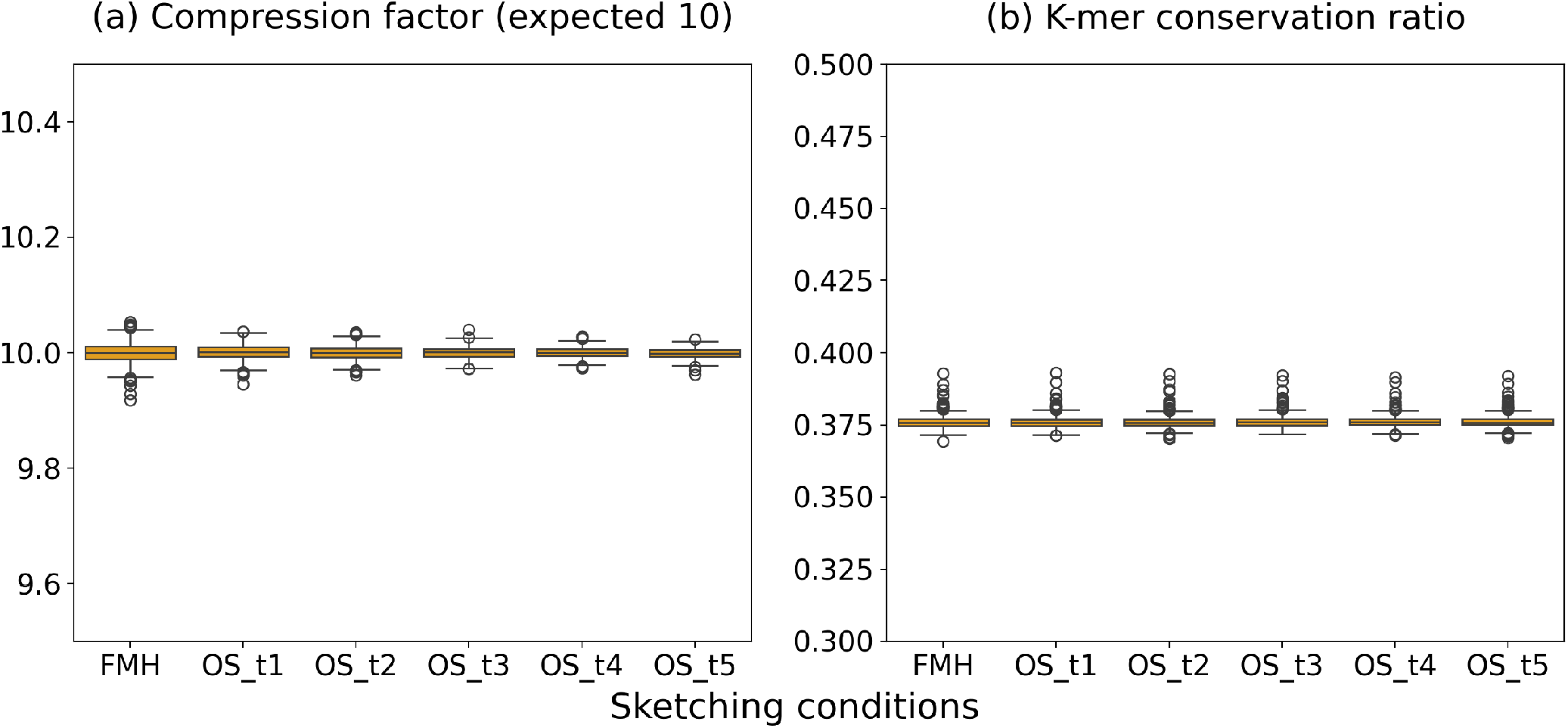
Open syncmer vs FracMinHash: compression ratio and *k*-mer conservation are consistent. Boxplot of compression ratio for the FracMinHash sketch with *c* = 10 and the open syncmer sketch with *k* = 20, *s* = 11 and *t* ranging in *t* = 1…5. By choosing appropriate parameters, both the FracMinHash sketch and the open syncmer sketch can achieve (a) the same expected sketch size and (b) comparable *k*-mer level conservation.

### Similarity estimation

Here we evaluate the performance of using open syncmers for Jaccard and containment estimation, as outlined in the previous section. Following the construction of FracMinHash and open syncmer sketches for all input genomes (with parameters: *k* = 20, *s* = 11, *t* = 5, *c* = 10), we estimated their pairwise similarities using open syncmers and FracMinHash sketches. Ground truth values were determined through brute-force calculations of all *k*-mers from the reference genomes. In theory, open syncmer sketches should produce identical results (Section 2. 2.3) and this is indeed what we observed in Figure 4.

**Fig. 4.**
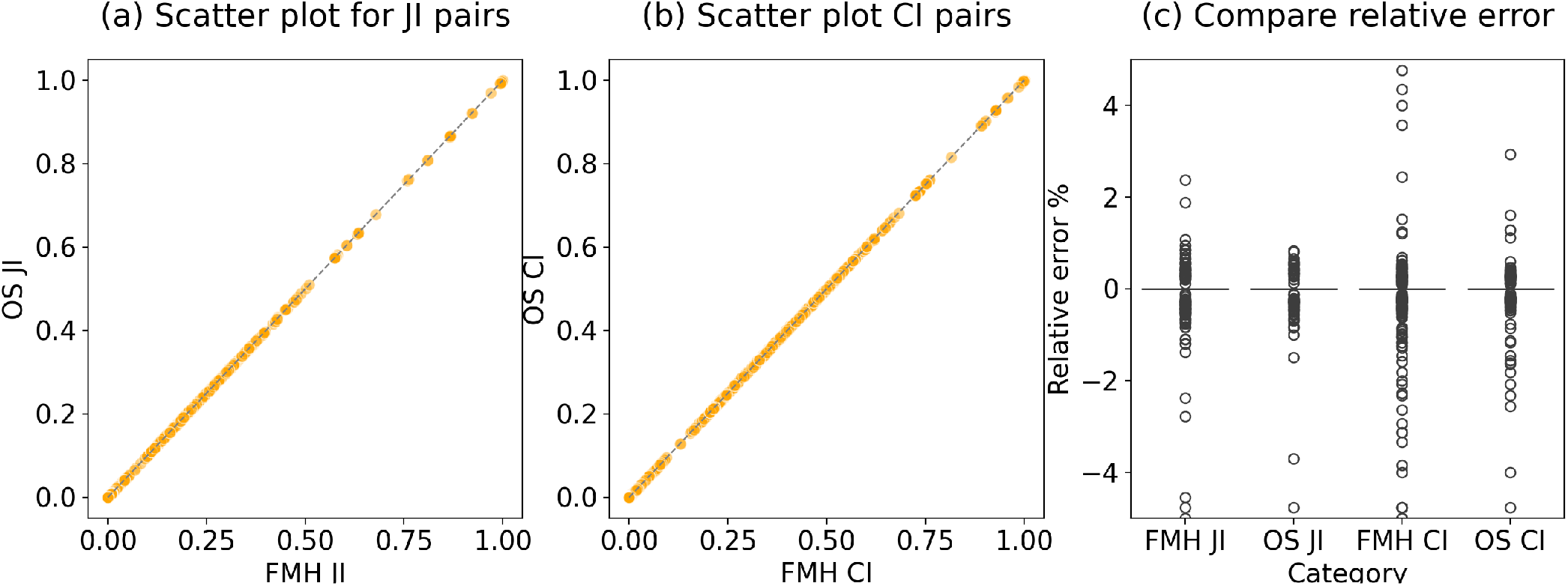
The open syncmer sketch fits the MinHash algorithm. (a) The scatter plot of Jaccard Index (JI) estimation by open syncmers and FracMinHash. The data points align closely along the *y* = *x* line, indicating a high degree of consistency between the two methods. (b) The scatter plot of containment index estimation by open syncmers and FracMinHash. (c) Relative error (in percentage) for all JI and CI estimations are small and comparable between the two methods.

**Fig. 5.**
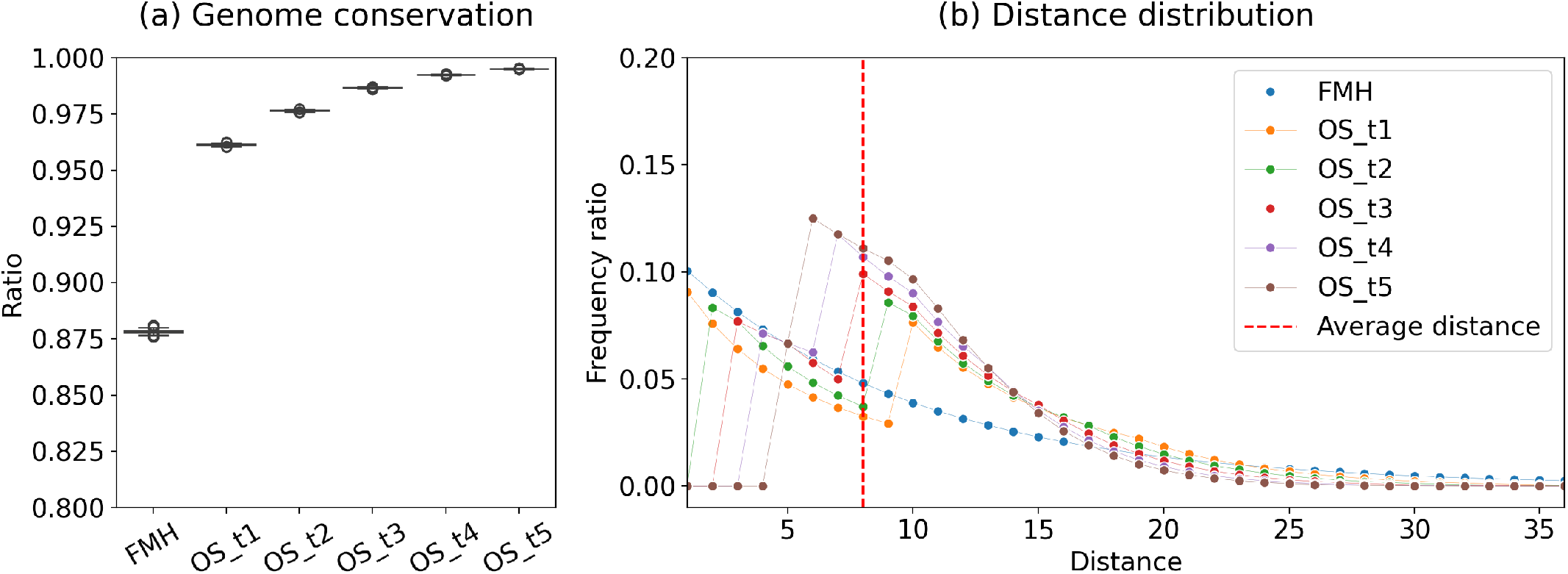
genome conservation and distribution of *k*-mer distances. (a) Boxplot of genome conservation by the FracMinHash sketch with *c* = 10 and the open syncmer sketch with *k* = 20, *s* = 11 and *t* ranging in *t* = 1…5. Open synmcers provide better genome conservation when the value of *t* increases. (b) Lineplot for the average distances between adjacent *k*-mers along the genome, under condition *k* = 12, *s* = 5. Distances are more centered around the mean when the value of *t* approaches its optimal location (OS t5). And we can observe the “window-free” event for different *t* values.

### Distance distribution

The results above aligns with our expectation that the open syncmer sketch is on par with the FracMinHash sketch when it comes to Jaccard and containment estimation (i.e., *k*-mer conservation between two sets). While the status of *k*-mer matching remains constant, it’s noteworthy that open syncmers offer a window-free guarantee, a situation where 2 consecutive *k*-mers in the sketch are at least *t* base apart until *t* reaches the midpoint of the *k*-mer.

This window-free feature, though not affecting *k*-mer conservation, will increase overall genome conservation. It has been proven by Shaw and Yu that the maximum genome conservation is achieved when *t* is the midpoint [20, Theorem 8]. In figure 5 we illustrated that genome conservations in all input genomes increase with increased *t* value (*k* = 12, *s* = 5); and the distance distribution will shift towards the mean to give a more evenly distributed coverage.

### Multi-resolution estimation and seed matching

With parameters (*k* = 20, *K* = 30, *s* = 11, *t* = 5), we created multi-resolution open syncmer (MOS) sketches for all the 420 random genomes and subsequently assessed *k*-mer conservation across various mutation rates for a spectrum of *k* values.

For each reference genome, we utilized a simple random point mutation model to generate ten mutated genomes. Subsequently, we generated Open Syncmer (OS) sketches and multi-resolution Open Syncmer (MOS) sketches with the same *k* value and employed them independently to compute the average *k*-mer conservation between the original genome and the mutated genomes.

Our observations indicate that the MOS sketches exhibit similar performance to regular open syncmers, implying their capacity to offer multi-resolution similarity estimation without the need to reiterate the sketching workflow with a different *k* size (Fig 6). This insight also presents an optimization strategy for addressing duplicate seed matching in read aligners: we can sketch with a larger *k* value and scale it down to accommodate an appropriate matching status; or we may expand hits with multiple matches to minimize duplication.

**Fig. 6.**
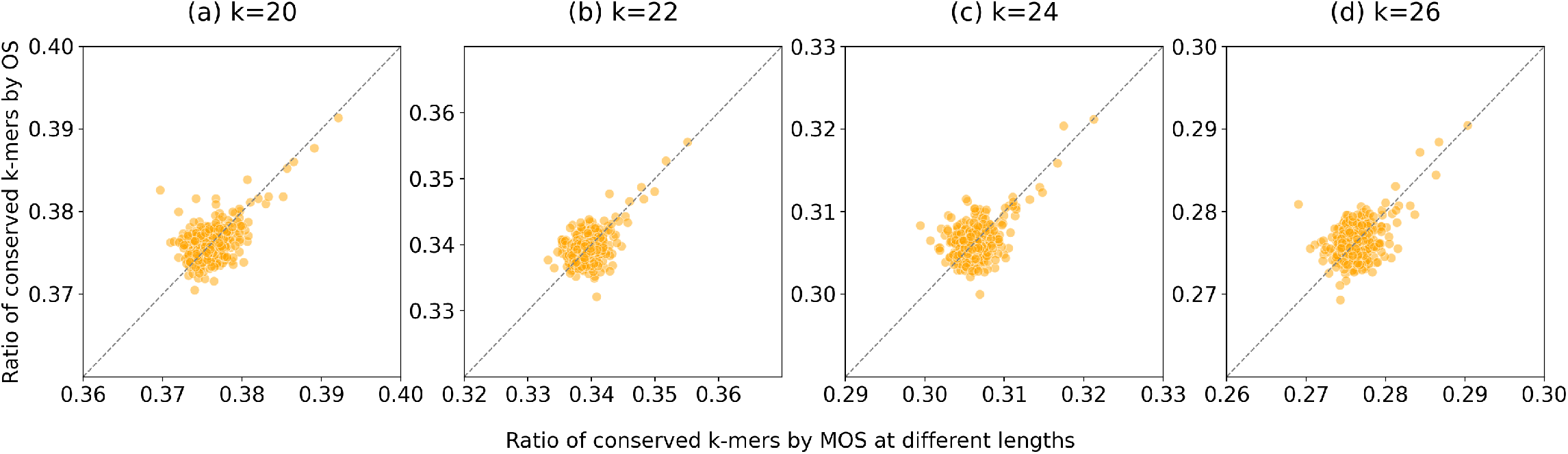
Multi-resolution open syncmers are comparable to regular open syncmers. We used the parameter *k* = 20, *K* = 30, *s* = 11, *t* = 5 to build the multi-resolution open syncmer sketches and compared their *k*-mer conservation to regular open syncmers at (a)*k* = 20, (b)*k* = 22, (c)*k*=24, and (d)*k*=26. In addition to the variability introduced by the random mutation process, the multi-resolution estimation aligns closely with regular open syncmers, highlighting the capacity of MOS to facilitate multi-resolution estimation.

## Conclusion

This article presents a comparison between two sketching methods: open syncmers and FracMinHash. Open syncmers have found successful applications in seed construction for read aligners, while the FracMinHash algorithm offers considerable power in metagenomic analysis with set similarity estimation. Our demonstration reveals that open syncmer sketches are equivalent to the FracMinHash sketch within the context of *k*-mer-based set similarity estimation, despite the fact that overlapping *k*-mers are not independent of being selected as open syncmers. However, the open syncmer sketch is still considered a random sample from the set of *k*-mers of a genome because all hash values are uniformly and randomly distributed (Section 2.2.3).

This dependence of overlapping *k*-mers, as observed in open syncmers, offers specific advantages. It allows us to establish a “window-free” guarantee, ensuring that the distance between consecutive *k*-mers has a lower bound. This leads to a more evenly distributed distance pattern (though still having the same average distance), providing higher genome conservation. While both possess comparable and linear computational costs, open syncmer sketches are advantages over regular FMH sketches: they are equivalent in terms of *k*-mer-based similarities but the former exhibits superior seeding performance to preserve genomic information. This attribute is particularly beneficial for seed matching in read alignment.

Furthermore, we have extended our previous work related to multi-resolution *k*-mer-based estimation to incorporate open syncmers, resulting in a novel sketching strategy for multi-resolution open syncmers (MOS). MOS sketches maintain the same capability to the regular open syncmer sketches for estimating *k*-mer conservations and can be employed for multi-resolution similarity estimation, as well as for seed matching in alignment tasks.

### Future direction

The analyses described above can be extended to other syncmer settings, such as closed syncmers [5] or PSS [4]. However, the performance of open syncmers is already satisfactory. The combination of being “window-free” with a large *t* value and having a constant expected size ensures that: most distances between adjacent *k*-mers will be smaller than *k*, resulting in a high genome conservation (Figure 5). The next step needed is comprehensive benchmarking analyses on metagenomics to showcase the comparable efficacy of OS and MOS in applications where FracMinHash is widely used; and then try to explore the potential advantages stemming from enhanced genome conservation.

One challenge when applying open syncmers in read alignment practice is the issue of duplication. Kristoffer tried to address this concern by utilizing open syncmers in combination with strobermers for seed construction [18]. Our work on MOS offers an alternative approach: since we can easily modify seed sizes without compromising performance (Figure 6), MOS could be integrated into the seed matching step of the read aligner, potentially mitigating the duplication problem.

## References

1. Austin Appleby. Smhasher. https://github.com/aappleby/smhasher, 2008.

2. Mahdi Belbasi, Antonio Blanca, Robert S Harris, David Koslicki, and Paul Medvedev. The minimizer jaccard estimator is biased and inconsistent. Bioinformatics, 38(Supplement 1):i169–i176, 2022.

3. Andrei Z Broder. On the resemblance and containment of documents .In Proceedings. Compression and Complexity of SEQUENCES 1997 (Cat. No. 97TB100171), pages 21–29. IEEE, 1997.

4. Abhinav Dutta, David Pellow, and Ron Shamir. Parameterized syncmer schemes improve long-read mapping. PLOS Computational Biology, 18(10):e1010638, 2022.

5. Robert Edgar. Syncmers are more sensitive than minimizers for selecting conserved k-mers in biological sequences. PeerJ, 9:e10805, 2021.

6. Bariş Ekim, Bonnie Berger, and Rayan Chikhi. Minimizer-space de bruijn graphs: Whole-genome assembly of long reads in minutes on a personal computer. Cell systems, 12(10):958– 968, 2021.

7. César Estébanez, Yago Saez, Gustavo Recio, and Pedro Isasi. Performance of the most common non-cryptographic hash functions. Software: Practice and Experience, 44(6):681–698, 2014.

8. Martin C Frith, Laurent Noé, and Gregory Kucherov. Minimally overlapping words for sequence similarity search. Bioinformatics, 36(22-23):5344–5350, 2020.

9. Mahmudur Rahman Hera, N Tessa Pierce-Ward, and David Koslicki. Deriving confidence intervals for mutation rates across a wide range of evolutionary distances using fracminhash. Genome Research, pages gr–277651, 2023.

10. Luiz Irber, Phillip T Brooks, Taylor Reiter, N Tessa Pierce-Ward, Mahmudur Rahman Hera, David Koslicki, and C Titus Brown. Lightweight compositional analysis of metagenomes with fracminhash and minimum metagenome covers. bioRxiv, pages 2022–01, 2022.

11. Chirag Jain, Luis M Rodriguez-R, Adam M Phillippy, Konstantinos T Konstantinidis, and Srinivas Aluru. High throughput ani analysis of 90k prokaryotic genomes reveals clear species boundaries. Nature communications, 9(1):5114, 2018.

12. David Koslicki, Stephen White, Chunyu Ma, and Alexei Novikov. Yacht: an ani-based statistical test to detect microbial presence/absence in a metagenomic sample. bioRxiv, pages 2023–04, 2023.

13. Heng Li. Minimap2: pairwise alignment for nucleotide sequences. Bioinformatics, 34(18):3094–3100, 2018.

14. Shaopeng Liu and David Koslicki. Cmash: fast, multi-resolution estimation of k-mer-based jaccard and containment indices. Bioinformatics, 38(Supplement 1):i28–i35, 2022.

15. Brian D Ondov, Gabriel J Starrett, Anna Sappington, Aleksandra Kostic, Sergey Koren, Christopher B Buck, and Adam M Phillippy. Mash screen: high-throughput sequence containment estimation for genome discovery. Genome biology, 20:1–13, 2019.

16. Donovan H Parks, Maria Chuvochina, Christian Rinke, Aaron J Mussig, Pierre-Alain Chaumeil, and Philip Hugenholtz. Gtdb: an ongoing census of bacterial and archaeal diversity through a phylogenetically consistent, rank normalized and complete genome-based taxonomy. Nucleic acids research, 50(D1):D785–D794, 2022.

17. N Tessa Pierce, Luiz Irber, Taylor Reiter, Phillip Brooks, and C Titus Brown. Large-scale sequence comparisons with sourmash. F1000Research, 8, 2019.

18. Kristoffer Sahlin. Strobealign: flexible seed size enables ultra-fast and accurate read alignment. Genome Biology, 23(1):260, 2022.

19. Kristoffer Sahlin, Thomas Baudeau, Bastien Cazaux, and Camille Marchet. A survey of mapping algorithms in the long-reads era. Genome Biology, 24(1):1–23, 2023.

20. Jim Shaw and Yun William Yu. Theory of local k-mer selection with applications to long-read alignment. Bioinformatics, 38(20):4659–4669, 2022.

21. Jim Shaw and Yun William Yu. Fast and robust metagenomic sequence comparison through sparse chaining with skani. bioRxiv, pages 2023–01, 2023.

22. Yoshihiro Shibuya, Djamal Belazzougui, and Gregory Kucherov. Efficient reconciliation of genomic datasets of high similarity .In 22nd International Workshop on Algorithms in Bioinformatics (WABI 2022), volume 242 of Leibniz International Proceedings in Informatics (LIPIcs), pages 14:1–14:14. Schloss Dagstuhl - Leibniz-Zentrum für Informatik, 2022.

23. Derrick E Wood and Steven L Salzberg. Kraken: ultrafast metagenomic sequence classification using exact alignments. Genome biology, 15(3):1–12, 2014.

